# Single-embryo metabolomics reveals developmental metabolism in the early *Drosophila embryo*

**DOI:** 10.1101/2024.04.17.589796

**Authors:** J. Eduardo Pérez-Mojica, Zachary B. Madaj, Christine N. Isaguirre, Joe Roy, Kin H. Lau, Ryan D. Sheldon, Adelheid Lempradl

## Abstract

Early embryonic development is characterized by the transition from maternal factor reliance to zygotic control. These processes set the stage for the embryo’s basic structure and cellular differentiation. While relatively detailed knowledge exists of the transcriptional events during early development, little is known about the concurrent metabolic processes. Understanding these processes, however, is important since they are linked to cell fate determination and organ and tissue formation. The primary reasons for the limited progress in the field are technical limitations due to the small amount of material available during early embryonic time windows. Here, we introduce a novel single-embryo methodology that places us in an exciting position to analyze the early embryo’s metabolome and transcriptome in an integrated manner and at high temporal resolution. The resulting data allow us to map concomitant metabolic and transcriptional programs in early *Drosophila* embryonic development. Our results reveal that a substantial number of metabolites exhibit dynamic patterns with some changing even before the onset of zygotic transcription. dNTPs for example show a temporal pattern that correlates with cell division patterns in the early embryo. In summary, here we present an operationally simple single-embryo metabolomics methodology and provide a detailed picture of early developmental metabolic processes at unprecedented temporal resolution.

## Main

In animal development the earliest stages are controlled by maternally deposited biomolecules, including mRNAs and metabolites. Because ultimately the maternal contributions will become insufficient to sustain development, the zygote needs to take control. This process is collectively referred to as maternal-to-zygotic (MZT) transition^1^. During this period the zygote initiates transcription of its own genome, degrades or recycles the factors deposited by the mother, and starts producing its own biomolecules. While processes such as zygotic genome activation (ZGA) have been studied extensively^2^, the metabolic details of early development have been largely inaccessible. Studying metabolism at early stages of development is challenging given the small amount of material available, the rapid progression of development, and, in mammals, the continuous nutritional inputs from the mother though the placenta. As a closed system (e.g., no further input from the mother after laying), oviparous animal models, such as *Drosophila*, overcome the last of these issues, allowing us to focus exclusively on understanding embryonic metabolic programs, without any confounding inputs from the mother. To overcome the challenge of limited material, prior studies in *Drosophila* have utilized pooled embryo samples from 2 or 4-h developmental windows and focused on the metabolite changes that occur over the entirety of embryogenesis^3,4^. However, to understand how the zygote takes control over its own metabolism and how zygotic metabolism integrates with transcriptomic programs, high-resolution data are required.

We present an operationally simple single-embryo transcriptomic and metabolomic methodology developed to observe early developmental metabolism in real time. The resulting dataset provides an unprecedented view of the dynamic processes underlying early developmental metabolism. We previously reported a single embryo RNA-sequencing (RNA-seq) approach to generate high-resolution transcriptome datasets from *Drosophila* embryos^5,6^. Here we have enhanced this method by utilizing genetically different fly strains from the *Drosophila Genetic Reference Panel* (DGRP) for allele specific expression analysis^7,8^. Furthermore, we have expanded our approach by integrating metabolomics into our methodology, enabling multi-’omics analysis^9^. Our novel approach allows us to characterize metabolite profiles and allele-specific expression during early embryonic development (0-3 h in *Drosophila*) (**Figure 1A)**.

**Figure 1.**
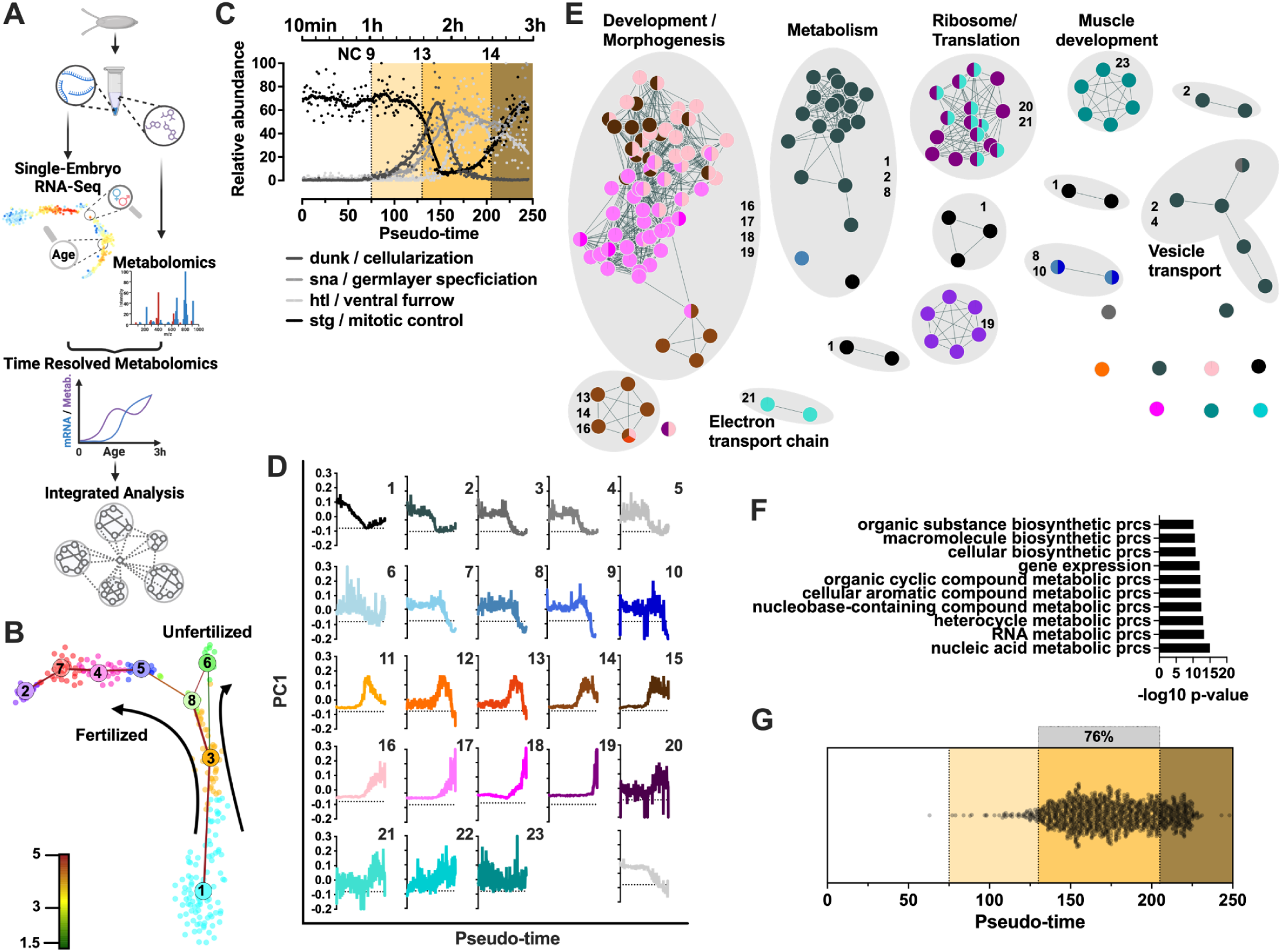
Transcriptional landscape of early Drosophila development. **(A)** Schematic representation of the method: single eggs are collected, and metabolites and RNA isolated from the same sample. Metabolites are used for metabolomics while RNA is processed for sequencing by a modified CEL-Seq2 protocol. RNA-seq data is analyzed to determine embryo age and sex (see methods for details). **(B)** t-SNE map visualization of RNA-seq of embryos (10min - 3h) and unfertilized eggs. k-medoids clusters are indicated by different colors and the intercluster links are indicated by straight lines with the color indicating the significance of the link, from red as the strongest to green as the weakest. **(C)** Normalized gene expression of dunk, disrupted underground network; sna, snail; htl, heartless; and stg, string plotted along our pseudo-time order. Estimated time and nuclear cycle (NC) are indicated at the top. **(D)** First principal component (PC1) or eigengene E of each module generated by WGCNA plotted along our pseudo-time order. Graphs represent the gene expression profile of all transcripts in a given module. **(E)** Network map of overrepresented pathways in modules, pathways are linked according to gene-set overlap. Overrepresentation analysis using g:profiler (significance threshold g:SCS=0.1) was used as input into Cytoscape for network visualization. **(F)** 10 most enriched pathways by adj. p-value, using g:profiler (significance threshold g:SCS=0.1) of zygotically expressed genes; prcs, process **(G)** Activation pseudo-time for each of the zygotically expressed genes, the gray bar indicates the NC 14 during which 76% of all genes start expression.

### Determining embryo age using a high-resolution transcriptome dataset

First, to determine the age for each individual embryo we generated a single-embryo transcriptome dataset using a modified version of our previously published approach^5^. Instead of directly isolating RNA, we first performed metabolite extraction using 80% methanol and isolated the RNA from the remaining pellet using TRIzol. Within our dataset we detected 7367 different transcripts with a minimum number of 5 unique reads in at least 5 individual embryos. The resulting time-resolved RNA-seq dataset includes the transcriptomes of 245 single embryos and 22 unfertilized eggs (>150K unique transcripts/embryo, average of 760K reads/embryo), a resolution of around 1.4 embryos per minute, twice that of our previous study^5^. **Figure 1B** shows a t-SNE map with the inferred differentiation trajectory indicated for embryos and unfertilized eggs. Ordering the embryos along this pseudo-temporal trajectory reaffirmed the dispensability of manual staging (**Figure S1A**) and the absence of batch effects for our protocol (**Figure S1B**). The transcriptome entropy of embryos along this developmental trajectory mirrored the loss in entropy seen for differentiation trajectories of single-cells^10^ (**Figure S1C**), further confirming the validity of our pseudo-temporal trajectory ( **Figure S1C**). An analysis of raw counts along the pseudo-time trajectory showed a change in the number of transcripts over time, with older embryos containing less transcripts (**Figure S1D**).

Since RaceID3 uses a correlation-based metric for the computation of the distance object, the determination of clusters and pseudo-time trajectory are independent of normalization^11^. Hence, we applied the Remove Unwanted Variation Using Control Genes (RUVg) procedure^12^ for data normalization and follow-up analysis. As the set of control genes for RUVg, we used the 10 least variable genes in each gene expression quartile from our previously published dataset^5^. Next, we determined the developmental stages reflected by our pseudo-temporal trajectory. To this end we plotted the dynamic expression patterns of genes regulating important developmental milestones of *Drosophila* embryogenesis, such as cellularization (*nullo*), germ layer specification (*snail*, *sna*), ventral furrow formation (*heartless*, *htl*) and mitotic control (*stinger*, *stg*)^13,14^ (**Figure 1C**). This analysis positions NC 1-8 prior to pseudo-time 75, NC 9-13 between pseudo-time 76 and 146, NC 14 between 147 and 201, and the cellular blastoderm thereafter. Using our previously validated approach^6^, we are also able to identify male and female embryos using their transcriptome for approximately half of the embryos (pseudo-time 127 onwards)(**Figure S1E, S1F, and Table S1**) and determine sex-specific differences on gene expression (**Table S2**). In summary, these results are evidence of the unprecedented temporal resolution achieved by using our single-embryo transcriptome methodology.

### Continuous transcriptome analysis reveals dedicated expression modules for metabolic pathways

Next, we wanted to see if certain pathways show coordinated patterns of expression during early development and how these relate to metabolism. To identify highly correlated genes across our time window, we used WGCNA (Weighted Gene Co-expression Network Analysis)^15,16^. After merging modules with high topological overlap, we identified 23 modules with more than 20 genes and distinct transcriptional patterns (**Figure 1D, S1G, and S1H**). Around 58% of all transcripts fall within modules with a pattern dominated by maternal deposition and zygotic degradation (modules 1 through 10). Only 23% show a pattern dominated by zygotic transcription. This dominance of maternal degradation could explain the reduced number of transcripts over time. The remaining 1418 transcripts out of 7367 did not fall within any of the identified modules **(Table S3)**. Using these modules we performed overrepresentation analysis of Gene Ontology (GO), KEGG and Wikipathway terms using g:profiler (significance threshold g:SCS=0.1)^17–21^. Modules 4, 5, 6, 7, 9, 11 and 12 showed no significant pathway enrichment. The enrichments for the remaining modules were used to create a network map using the Cytoscape Compound Spring Embedder (CoSE) layout function. Nodes were color-coded based on the module in which they are enriched **(Figure 1E and S1I)**.

The resulting map reveals several sub-networks of related pathways dominated by specific modules. A completely annotated network and a list of pathway enrichments can be found in **Figure S1I** and **Table S4**. A subset is discussed here and displayed in **(Figure 1E).** Specifically, there is a vast web of development/cell fate commitment related pathway terms driven by modules comprised of 1175 genes which display transcription patterns dominated by zygotic gene expression (modules 14, 15, 16, 17 and 18, **Figure 1D**). Module 23 which is enriched in muscle development pathways shows variable but overall stable expression. The remaining modules show enrichments in pathways that are not classically associated with development (**Figure 1E** and **S1I**). This includes modules which are enriched in pathways related to metabolism and exhibit distinct patterns of maternal degradation (modules 1, 2, 8). Interestingly, module 2, which is enriched for various lipid metabolic pathways also shows enrichment for vesicle transport, supporting the recently reported importance of vesicles in nutrient sorting in the early embryo^22^. Seemingly uncoupled from this general pattern of metabolic pathways are electron transport chain pathway related genes which are enriched in module 21 and are showing a highly variable pattern characterized by zygotic expression. This indicates that the transcription of energy metabolism related genes is uncoupled from those involved in biosynthetic pathways. The same general pattern observed for genes related to the electron transport chain is also evident in ribosome/translation pathways (modules 21 and 22). In summary, our analysis reveals dedicated modules of transcription with distinct patterns of expression for genes involved in developmental and metabolic pathways.

### Revealing the exact onset of transcription using allele specific analysis

While the transcriptional modules in **Figure 1D** reflect the patterns of transcript abundance, widespread maternal mRNA decay in early embryos masks the true onset of gene transcription. To enable the assessment of the accurate transcriptional activation for genes, we used two genetically different DGRP lines (males/DGRP_352, females/DGRP_737) with known genetic variations in our crosses^7,8^. Variant calling on the RNA-seq data derived from hybrid offspring was carried out by a modified GATK’s workflow retaining only known and biallelic SNPs annotated to a single gene^7^. We detected a total of 2453 genes with SNPs in our dataset. Since unfertilized eggs should only contain maternal transcripts but none from the paternal allele, we excluded all transcripts with paternal reads present in unfertilized eggs (≥2 in any unfertilized egg, n=395) from downstream analysis.

Upon fertilization, *Drosophila* embryos initially contain trace amounts of paternal mRNAs delivered by the sperm. It is only when the zygote assumes control over its genome that transcripts from both parental alleles become detectable. We thus used the paternal allele data to identify transcripts that are zygotically expressed during early development. This analysis revealed that around 60% of genes (n=1476) with SNPs are expressed from the paternal allele (minimum 3 paternal transcripts in at least 10 embryos) at some time during our 3h window, this agrees with a previous report, which reported about 44% of genes as non-expressed during early development^23^. Overrepresentation analysis showed that zygotically expressed genes are enriched in pathways related to transcription, with the nucleic acid metabolic process as the most enriched pathway (**Figure 1F**). To establish the exact onset of transcription, generalized additive models were used to estimate, with 95% confidence, the earliest pseudo-time where the mean expression was at least one unique read. This analysis revealed that transcription of 76% of all zygotically transcribed genes (n=1126 is initiated during cycle 14 within a window of 75 points in the pseudo-time (from 130 to 204), which translates to an approximate 50 min time window **(Figure 1G, Table S5).** These findings are consistent with prior studies that assessed nascent zygotic mRNAs^24^. Significantly, our pseudo-time methodology accurately determines embryo age, thus circumventing previously reported issues of mRNA contamination from older embryos during allele-specific analysis of zygotic genome activation^24,25^. Classification of these genes into activation time windows revealed 4 major waves of gene activation (**Figure S1J**), with the 2^nd^ wave representing transcriptional activation during cycle 14. Collectively, utilizing genetically diverse *Drosophila* strains in our crosses allows for allele specific analysis providing a high-resolution picture of zygotic genome activation in *Drosophila*.

### Single-embryo metabolomics reveals time-resolved metabolite profiles

While our transcriptome analysis is detailed in nature, it offers limited insights into metabolic processes during development. To gain additional insights into early embryo metabolism, we combined our high-temporal resolution transcriptional analysis with single-embryo metabolite profiling. Here, we present this novel methodology and reveal unprecedented insights into the molecular metabolic mechanisms underlying early development. Because individual *Drosophila* embryos are small, metabolite abundances are therefore low. To narrow down the analysis on a subset of polar metabolites, we performed LC-MS on metabolite extracts with concentrations equivalent to 10, 5 and 1 embryo. This resulted in a panel of 155 metabolites in our final analysis. To assess metabolism during early development in a time dependent manner, we targeted this panel of metabolites on metabolite extracts isolated from the same samples included in our pseudo-time analysis (**Figure 1B**, 245 single embryos and 22 unfertilized eggs). Previous work from our group demonstrates the phenotypic veracity of this single-sample multi-’omic workflow^9^. **Figure 2A** shows a schematic for the metabolite data processing.

**Figure 2.**
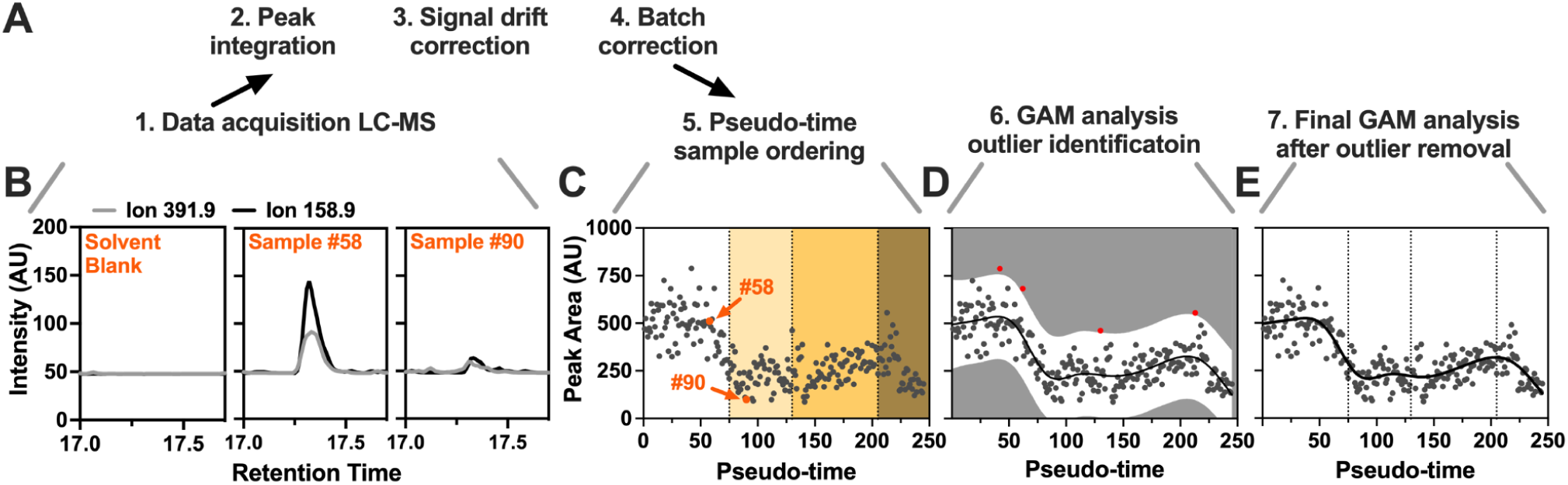
Workflow for single-embryo time-resolved metabolomics. **(A)** Method overview. **(B)** Ion chromatograms for two transitions (490➝391.9, 490➝158.9) of dATP for a solvent blank, sample #58, and sample #90. **(C)** dATP abundance in arbitrary units (AU) plotted along the RNA-seq pseudo-time order. Arrows indicate samples #58 and #90 plotted in B, with high or low relative dATP abundance, respectively. **(D)** A generalized additive model (GAM) using the pseudo-time aligned metabolite data identifies outliers (red data points) based on 99.7% prediction intervals (white area). **(E)** After outlier removal, GAM is applied again to determine the predicted values for each metabolite (black line).

Despite the low sample concentration in our single embryo samples, we detected unambiguous signals for 81 metabolites (**Figure S2A, Table S6 and S7**). As an example, **Figure 2B** shows extracted ion chromatograms for two transitions (490➝391.9, 490➝158.9) of dATP from two representative embryo samples chosen from two distinct pseudo-times. Inclusion of solvent blank injection demonstrates clear signal in the biological samples. Following data acquisition and initial processing (see methods), samples were arranged along the pseudo-time trajectory derived from our transcriptomics analysis to reveal the dynamic behavior of individual metabolites during early development. **Figure 2C** shows the metabolomics results for dATP presented along the pseudo-time trajectory. Despite low input and relatively low dATP signal abundance, we observed a complex pseudo-time dependent response. Importantly, these dynamic changes in dATPs levels were devoid of batch effects (**Figure S2B**) and were not observed in unfertilized eggs (**Figure S2C**). To navigate the inherent noise due to analytical and technical variation, especially given the single-embryo focus, we employed data fitting and smoothing through Generalized Additive Models (GAM, **Figure 2D**). Sample outliers for each metabolite were identified using a conservative prediction interval of 99.7% and subsequently excluded from analysis. Following outlier removal, the GAM was re-run to establish the definitive pseudo-time pattern for each specific metabolite (**Figure 2E**). To accommodate the differences in relative abundance and enhance the clarity of visual data representation, graphs will show scaled data from here onwards (**Table S8**). Focusing on dATP, our analysis shows initial high levels of free dATP followed by a drastic drop at around NC 8 after which abundance remains constant for a short period of time (**Figure 2E**). This time window coincides with a time of increased nucleotide demand due to genome duplication during rapid nuclear division. During early embryogenesis, the first 8 cell cycles last only 8 min and progressively slow down to 18 min by cycle 13^26^. The dATP signal increases again during NC14, which reflects a time absent of nuclear divisions. Once the embryo resumes nuclear divisions after NC14, dATP abundance drops again. In summary, our methodology successfully detects 81 individual metabolites in single embryo samples (**Table S8**), providing an unprecedented view of early embryo metabolism. Utilizing the previously determined transcriptome pseudo-time we can attribute metabolite abundance to individual embryos. This approach allows us to observe early developmental metabolism in a continuous manner and provides a detailed understanding of the dynamic metabolic processes underlying embryonic development. Moreover, the dynamic pattern of dATP abundance aligns with the time course of nuclear divisions during early development.

### Revealing known and novel patterns of metabolite abundance

While our understanding of the dynamics of metabolism during early development remains limited, emphasis in *Drosophila* has been placed on deoxynucleotides (dNTPs). Previous studies have demonstrated fluctuations in dNTP levels during the initial stages of development, providing valuable insights into the metabolic dynamics associated with DNA synthesis^27,28^. To verify our results and demonstrate their biological significance, we focused our initial analysis on dNTPs temporal patterns. Our time resolved metabolite profiling shows the dynamic patterns in dNTP abundance during the first 3h of development. Both dTTP and dATP exhibit a sharp decline as early as pseudo-time 50, followed by an increase in abundance around pseudo-time 150 (**Figure 3A,D)**. dCTP, while showing a similar trend, displays a much more gradual decrease without a measurable rise at later time points (**Figure 3G**). Detection of dGTP is not possible by our method, which is likely attributed to a combination of low endogenous abundance and compound-intrinsic differences in ionization efficiency. Importantly, we validated all dNTPs signals with reference standards (**Figure S3A**).

**Figure 3.**
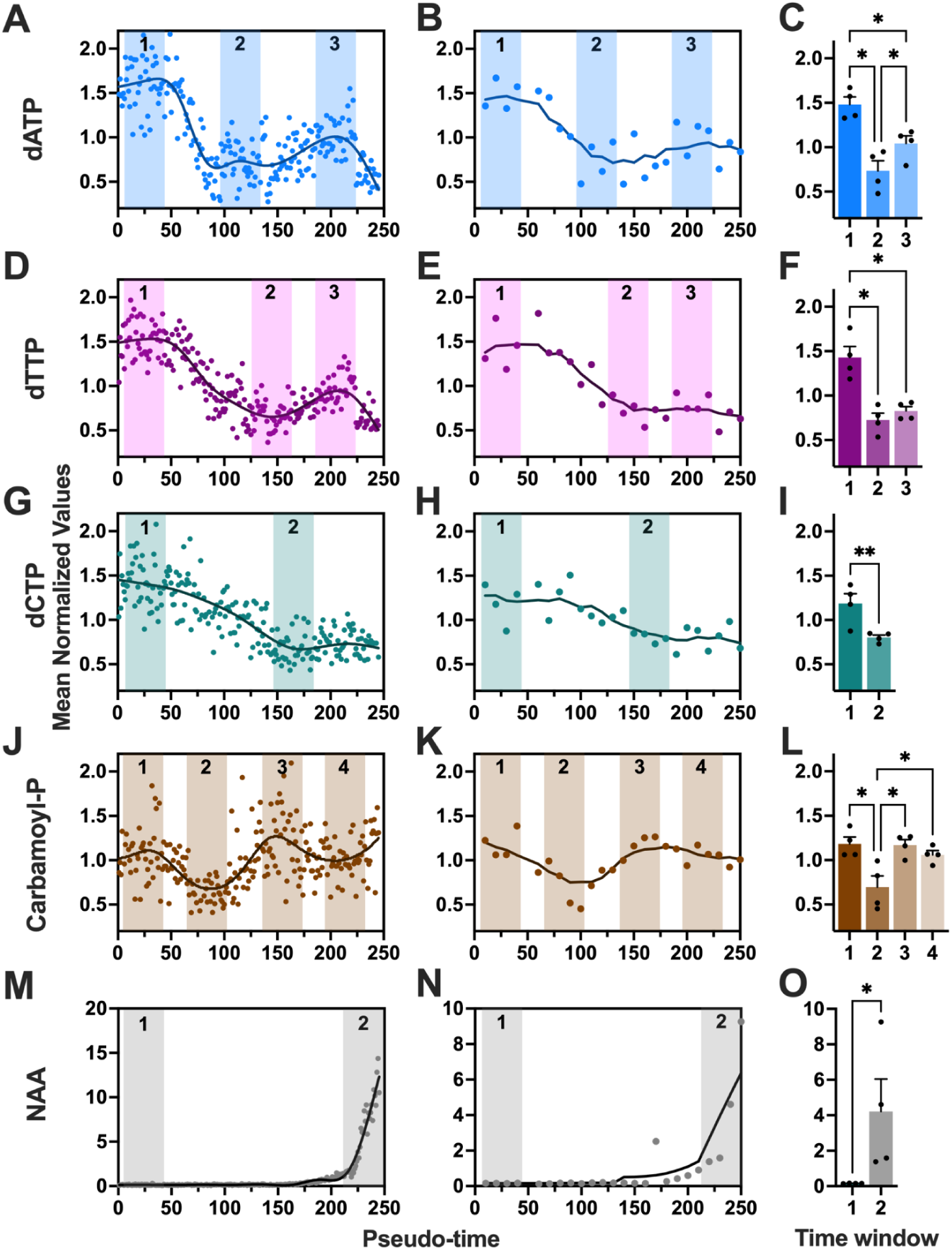
Single-embryo metabolite patterns reveal accurate patterns of metabolite abundance. **(A, D, G, J, M)** Metabolite relative abundance from single embryos plotted along the pseudo-time order. **(B, E, H, K, N)** Metabolite relative abundance from pooled embryos (n=10/sample) plotted along the pseudo-time order. The location of minima and maxima in the pseudo-time order is indicated by numbered colored areas. **(C, F, I, L, O)** Selected pooled samples for comparison (n=4 per group) based on abundance minima and maxima of the single-embryo pseudo-time order. Group means were compared (C, F, L) by one-way ANOVA or (I, O) by one-tailed unpaired t-test. Means that differ significantly are indicated by *, P < 0.05; **, P < 0.01.

Given the low metabolite signal in our single-embryo samples, we sought additional validation for the observed changes in samples with higher metabolite concentrations. For this purpose, we performed LC-MS on pools of 10 embryos to boost LC-MS signal intensity and minimize technical variation due to low-input. Since analysis of metabolites in male and female embryos did not reveal any significant sex-specific differences **(Figure S3B)** we disregarded sex for sample pooling. To preserve apparent developmental time, embryo metabolite extracts were pooled based on their RNA-seq pseudo-time, dried, resuspended and analyzed. Indeed, pooled samples demonstrated similar trends in dNTPs as those from single embryos (**Figures 3B, E, H**), therefore, supporting our single-embryo approach. To further validate the significance of the changes detected in our single-embryo dataset, we determined the abundance minima and maxima of our single-embryo smoothed temporal trajectory (shaded areas in **Figure 3A, D, G)** and assigned 4 consecutive pooled samples to the respective time windows. The results demonstrate that the initial decline in metabolite abundance is statistically significant for all three dNTPs (**Figures 3C, F, I)**. Importantly, our results from both, single- and pooled embryo samples, accurately mirror published metabolomics results from hand staged pooled embryos at defined developmental stages, confirming that dNTP’s are deposited by the mother and rapidly depleted during the initial 13 rounds of cell divisions^27,28^. Our data also reveal that at the onset of cellularization, when cell divisions are paused for approximately 30 min^29^, dATP and dTTP levels stabilize and even start to increase. While this increase is evident for both dATP and dTTP in our single-embryo time course, it is only significant for dATP using our pooled sample analysis. Surprisingly, despite the major transcriptional events taking place, ribonucleoside (NTP) levels were relatively stable during this 3h time window (**Figure S3C-J**). In summary, our single-embryo metabolomics methodology delivers a robust picture of nucleotide dynamics during early embryogenesis at unmatched resolution.

In addition to replicating known patterns of metabolites during early embryogenesis, our data also provide novel insights into 78 additional metabolites (**Table S8**). For example, carbamoyl-aspartate, which plays a crucial role in the urea cycle and pyrimidine biosynthesis, exhibits an oscillating pattern (**Figure 3J**). This oscillating pattern is reproduced in the pooled samples at a lower resolution (**Figure 3K)** and statistical analysis of pooled samples at minima and maxima shows significance for the first two comparisons (**Figure 3L)**. As another example, N-Acetyl Aspartate (NAA), which is predominantly found in neurons^30^, did not appear until later embryogenesis. Remarkably, our data show that NAA is produced *de novo* during NC14, before mature neurons develop (**Figure 3M)**. Like for the other metabolites, the observed pattern in our single-embryo data analysis is also captured in the pooled samples (**Figure 3N)** showing statistical significance (**Figure 3O).** Collectively, our data replicate known and reveal novel patterns of metabolite abundances. Using pooled samples at increased concentrations, we demonstrate that our single-embryo data accurately reflect dynamic processes during early embryonic development.

### Metabolite patterns are related to gene transcription

To determine if associations exist between metabolite levels and patterns of gene expression, we next evaluated correlations between WGCNA transcript modules and metabolite abundances. This analysis revealed extensive correlations among transcriptional modules (colored by module), while most metabolites (white circles) showed no significant correlation with any of the WGCNA modules (FDR < 0.05 and Pearson R^2 > 0.25)(**Figure S4A)**. The metabolites exhibiting significant correlations with transcriptional modules are shown in **Figure 4A** and include several amino acids, nucleotide related metabolites, NAA and GlcNAc-1-P. Given the overall absence of correlations between metabolites and transcript modules, we hypothesize that these few metabolites are central to embryonic development. To test this, we focused on dNTPs and its network neighbors. Analysis revealed direct positive correlations (red connecting lines) between dNTPs and modules 1 and 2, and negative correlations (blue connecting lines) with deoxyinosine, asparagine, and module 11 (**Figure 4B)**. Over representation analysis for modules 1 and 2 highlighted DNA replication and glycero- and phospholipid metabolic processes as the top enriched pathways (**Figure 4C and D**). Glycerophospholipids are the major lipid parts of cell membranes.

**Figure 4.**
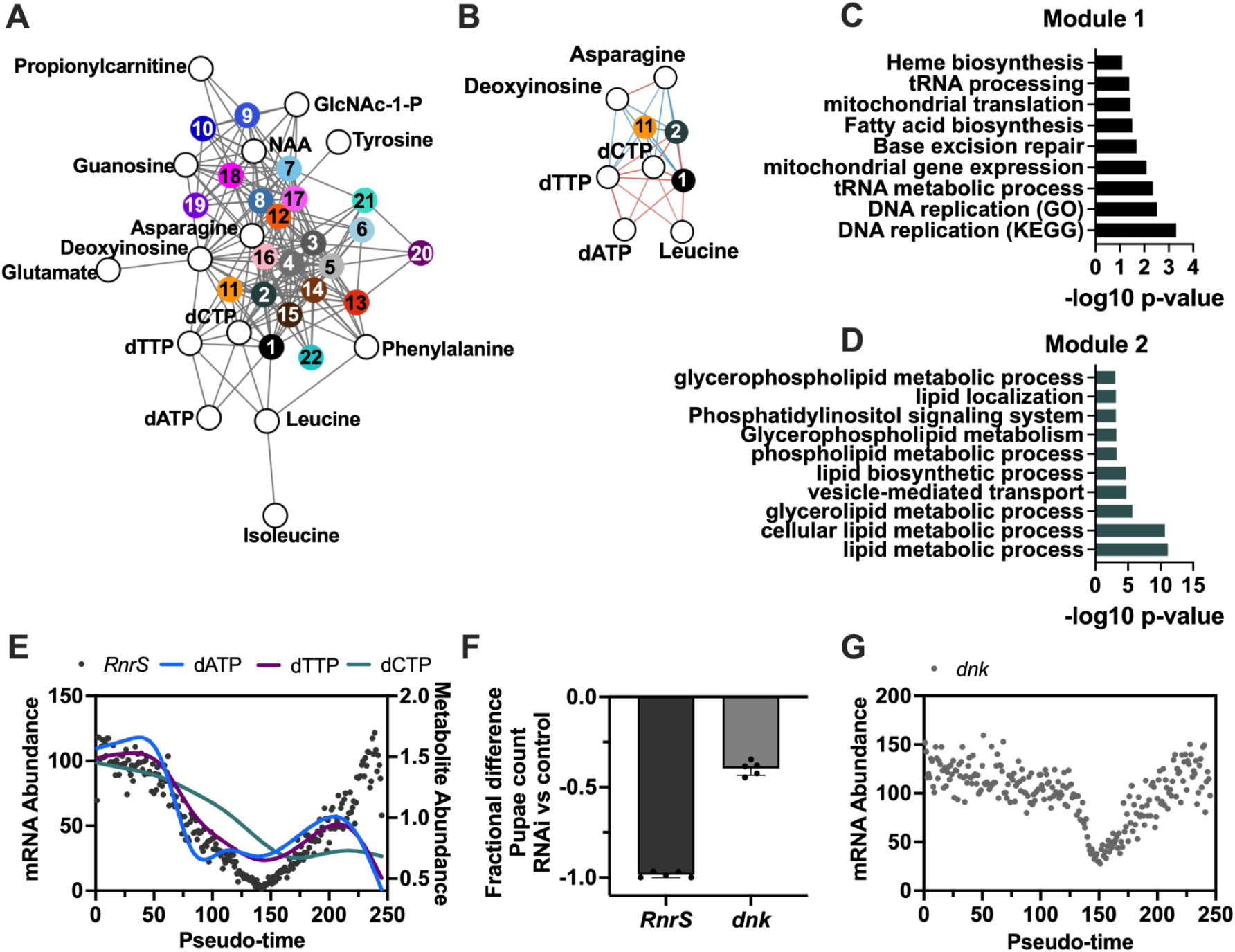
The interrelationship between metabolisms and transcription during Drosophila pre-gastrulation. **(A)** Network map of correlations between WCGNA transcript modules (colored circles) and metabolites (white circles) using network visualization with Cytoscape. **(B)** Network map of dNTPs and its neighbors. Red or blue connecting lines indicate positive or negative correlations, respectively. **(C, D)** Overrepresentation analysis for transcript in modules (C) 1 or (D) 2 using g:profiler (significance threshold g:SCS=0.1). **(E)** Normalized gene expression of RnrS, ribonucleoside diphosphate reductase small subunit (closed circles); and metabolite relative abundance of dNTPs (lines) plotted along our pseudo-time order. **(F)** Fractional difference in pupae count after oocyte-specific depletion of RnrS or dnk (deoxyribonucleoside kinase) by RNAi. **(G)** Normalized gene expression of dnk plotted along our pseudo-time order.

Notably, the rate limiting enzymes for de-novo deoxynucleotide synthesis, ribonucleotide reductases *RnrS* and *RnrL*, are members of the positively correlated modules 1 and 2 respectively **(Table S3)**. A closer examination of our single-embryo data revealed that an initial drop in dNTPs abundance coincides with the degradation of the maternally deposited *RnrS* transcript, which reaches its minimum at the onset of cellularization (pseudo-time 150, **Figure 4E**). Following this decline, *RnrS* exhibits zygotic expression after only a brief pause, coinciding with an increase in dATP abundance. These data suggest *RnrS* regulation by transcript abundance in addition to allosteric control by dNTPs^31^ (reviewed in a development context in ref^32^). A previous study using inhibitors showed that RNR activity in the early embryo is necessary for proper development despite the maternal deposition of dNTPs into the egg^27,28^. Using oocyte specific depletion of *RnrS* transcript^33^ we show that depletion of the maternally deposited transcript is sufficient to abrogate development. This result confirms the requirement for maternally deposited RNR (**Figure 4F**). Additionally, our data reveal that deoxyribonucleoside kinase (*dnk*), the rate limiting enzyme for dNTP salvage, is maternally deposited and follows a temporal pattern similar to *RrnS*, albeit shifted to later time points (**Figure 4G**). While no significant contributions from the salvage pathway were previously identified^28^, these results indicate that the salvage pathway’s contribution to the dNTP pool during development might have been underestimated. To confirm that this is indeed the case we depleted *dnk* transcripts in oocytes. In support of a significant contribution from the salvage pathway, depletion of *dnk* led to decreased offspring survival, but at a lower level than RNR knockdown (**Figure 4F**).

Another example of metabolites significantly correlated with WGCNA transcript modules is NAA **(Figure S4B)**. This nervous system-specific metabolite is positively correlated with modules (16, 17, 18, and 19), which are enriched in pathways associated with neuron development, as well as pentose and glucuronate interconversions (**Figure S4C-E**). In conclusion, our integrated analysis of time-resolved transcriptomics and metabolomics data in early *Drosophila* embryos reveals correlations between transcript modules and only a subset of metabolites. One of these relationships is between abundance of dNTPs and transcripts for genes involved in their biosynthesis. Crucially, we validate the significance of this relationship through maternal knockdown experiments. This, along with the findings related to NAA, support our hypothesis that nodes within our correlation network likely represent key processes central to embryo development.

## DISCUSSION

To gain comprehensive insights into early development processes has been challenging due to the limited amount of material, the confounding effects of pregnancy or embryo culture conditions, and the rapid progression of early embryogenesis. In this present study, we present an innovative methodology for analyzing dynamic metabolic processes during early embryonic development (0-3h in *Drosophila*). We build upon our previously published single-embryo transcriptomics approach, which uses the transcriptome to determine the developmental stage of the embryo^5,6^. We enhance this methodology in two key aspects. Firstly, we use genetically distinct *Drosophila* strains enabling the allele specific analysis of our transcriptome data. Secondly, and most significantly, we integrate the isolation of polar metabolites into our workflow to perform mass spectrometric analysis. These advancements enable, for the first time, assessment of the metabolite abundances in single embryos throughout the time-course of early embryonic development.

To develop our time-resolved single embryo multi-‘omics analysis we had to overcome two principal analytical challenges. First, due to rapid development and practical limitations defining the moment of fertilization, using traditional time-based collection for embryo collection lacked necessary developmental stage precision for informative metabolomics. To address this, we utilized single embryo RNA-seq signatures to accurately define developmental stage. However, this also necessitated the sequential extraction – and preservation – of both the metabolome and RNA of hundreds of embryos. We have previously demonstrated that RNA-seq following metabolite extraction with 80% methanol fully captures the phenotype detected by samples extracted for RNA-seq alone^9^. We employed this approach here, where samples were collected and homogenized in 80% methanol, and RNA-seq was performed on the insoluble fraction to accurately stage each embryo. Then, this RNA-based staging was used to place single embryo metabolomes across the developmental trajectory. Thus, we are able to construct a highly detailed metabolomic map of embryonic development.

The second analytical challenge was the small size (∼0.15 mm diameter) limiting the amount of metabolites available for metabolomics analysis. Unlike sequencing ‘omics approaches, the metabolome cannot be amplified to increase detection limits. Moreover, the metabolome is highly diverse and dynamic; it contains compounds with a wide range of chemical properties and concentrations, and structures. As such, no single LC-MS method can capture the entire metabolome. Thus, we turned to two orthogonal chromatography methods (reversed phase ESI+ mode and ion-paired negative mode) to maximize compound coverage. These were coupled to a triple quadrupole mass spectrometer, which excels at high sensitivity and specificity through MRM based transition monitoring, for analyte detection. Early method development efforts in pooled embryo samples allowed us to constrain the analytical scope to only compounds known to be present in these samples. This restricted our analysis only to useful transitions, in turn increasing instrument dwell-time per compound and, ultimately, further improving sensitivity. The power of this approach is evident in the current work, where we provide quantitative metabolomic coverage in high resolution across the developmental landscape. The power of this approach is apparent in the insight gained in deoxynucleotide metabolism. The demand for deoxynucleotides during the first stages of embryonic development is exceptionally high due to rapid cell divisions. This demand is only partially met through maternal deposition of dNTPs and *de novo* synthesis is required to sustain cell cycles after cycle 10^28^. Importantly, both too high and low dNTP levels in embryos have been shown to negatively impact development^27,28^, highlighting the critical need for precise control of dNTP levels. We uncover temporal associations that are indicative of the concurrent biological processes taking place during early embryonic development. These include rapid cell divisions during the initial cell cycles, a temporary cessation of cell division during nuclear cycle 14, and the reinitiation of cell divisions thereafter. Our network analysis reveals associations between the highly dynamic pattern for dNTPs and specific transcripts modules involved in cell cycle and membrane synthesis, suggesting an interdependency of dNTP synthesis with cell cycle and membrane synthesis for cellularization during early development. Notably, while maternal deposition of nucleosides and *de novo* synthesis have been recognized as the primary sources fulfilling the initial demand for dNTPs, our findings introduce the salvage pathway as an additional essential contributor to proper embryonic development.

Variability in sample collection, handling, extraction, and derivatization can affect the accuracy and reproducibility of metabolite measurements^34–36^. Here, we lysed intact embryos in the metabolite extraction solvent as the first step to ensure a stable extraction. Standardizing metabolomics procedures across different samples and conditions can be challenging, especially when dealing with large sample cohorts. However, in the case of *Drosophila* embryos, which function as a closed system and do not require an *ex vivo* culture step, their metabolite composition remains unaffected by harvesting methods. Leveraging this inherent strength of the model system, we can minimize technical noise and ensure robust data quality. Moreover, the fecundity of *Drosophila* allows for large sample sizes, enabling the detection of biologically relevant differences despite potential variability.

Analyzing multi-‘omics datasets requires sophisticated statistical and computational approaches to distinguish true biological signals from technical noise. To simplify the complexity inherent to such types of data, various data reduction methods including nonnegative matrix factorization, principal component analysis, and weighted gene co-expression network analysis (WGCNA) are commonly used. In our study we opted for WGCNA, a method that groups genes into modules based on similar expression patterns, in our context, analogous developmental patterns. Utilizing the first principal component of each module, known as the eigengene, as a representative transcription signal significantly reduced the number of gene-metabolite correlations from 7367*81 to 23*81. This not only enhanced computationally efficiency but also greatly reduced the multiple testing penalty. Moreover, by performing overrepresentation analysis on those modules, we gained valuable insights into the biological processes underlying significant correlations. Rather than focusing on individual gene-metabolite interactions, this approach enabled us to elucidate associations between metabolites and entire biological processes or molecular functions. The choice of generalized additive models (GAM) for regressing pseudo-time was motivated by the complexity of transcript and metabolic patterns in the first 3 hours of embryonic development. Given the complexity of these patterns, we required a flexible, data-driven method capable of accurately capturing sudden changes and identifying relevant maxima and minima along the time course. Leveraging the relatively large sample sizes using *Drosophila*, our GAM fits accurately capture the entire time course. Our analysis approach provides a blueprint for the future of single-cell methodologies, particularly as technology continues to advance. Similar to our dataset, these datasets will be large and complex, containing thousands of peaks or signals corresponding to different metabolites and transcripts.

In summary, our single-embryo methodology offers an unprecedented dataset with great value for the developmental biology community. By providing a detailed snapshot of molecular events at the individual embryo level, our approach unlocks new avenues for understanding the intricate processes governing embryonic development. Moreover, it presents a novel opportunity to investigate the impact of developmental perturbations on metabolic pathways, shedding light on the interplay between genetic and environmental factors during crucial stages of embryogenesis. This comprehensive dataset not only enriches our understanding of fundamental biological processes but also opens doors for future research aimed at unraveling the complexities of developmental metabolism and its implications for organismal health and disease.

## METHODS

### Fly stocks and embryo collection

Drosophila genetic reference panel (DGRP) 737 and 352 lines^5,6^ from Bloomington Stock Center (#83729 and #83728, respectively) were kept in incubators at 25°C with 60% humidity and a 12-hour light-dark cycle. All flies were raised at constant densities on standardized cornmeal food (Bloomington recipe), Fly food M (LabExpress, Michigan, USA), and DGRP_737 virgin females and DGRP_352 males transferred into 3 different embryo collection cages for 60 mm petri dishes (Genesee Scientific, USA) 3-4 days after eclosion. Quality control (QC) samples for metabolomics analysis (0-3 h post fertilization embryos) were collected a day before embryo collection started on 8-9 days old flies. An additional cage with only DGRP_737 virgin females was used for unfertilized eggs collection. In all experiments, food plates were changed and discarded twice before embryo or unfertilized egg collection started. For each sample collection, food plates were changed in time intervals to get 0-1 h samples and processed immediately or incubated for 1 or 2 more hours at 25 °C (1-2 h or 2-3 h samples, respectively). Samples were transferred into a pluriStrainer® 150 µM cell strainer (pluriSelect, USA) and washed with tap water, dechorionated by incubation in 3% sodium hypochlorite (PURE BRIGHT® bleach, KIK international LLC) for 4 min, washed in 120 mM NaCl (Sigma-Aldrich, USA) with 0.03% Triton X-100 (Fisher Scientific, USA) solution, and finally washed in ultrapure water (PURELAB® Ultra, ELGA). Single embryos or unfertilized eggs were transferred into 2 ml screw-cap tubes pre-filled with 1.4 mm ceramic beads (OMNI international, USA) using a 20/0 liner brush (Royal & Langnickel®, USA), snap-frozen on dry ice, and stored at -80°C.

### Metabolite extraction

Frozen embryos or unfertilized eggs were homogenized in 1 mL 80% methanol (v/v) (Optima™ LC/MS Grade, Fisher Chemical™) by bead-beating at 6 m/s for 30 seconds at 4°C using a Bead Ruptor Elite homogenizer coupled with Bead Ruptor Cryo Cooling Unit (OMNI international, USA). Samples were centrifuged for 10 minutes at 17,000 × g at 4°C and 800 µL of the soluble metabolite fraction transferred to new 1.5 mL tubes, dried (∼3 h) in a Genevac EZ-2 series evaporator (ATS Life Sciences Scientific Products) using low-BP lamp-off settings, and stored at -80°C. A similar process was carried out for the QC samples except that soluble metabolite fractions were combined into a single sample (n=300 embryos), mixed by vortex for 15 seconds, and a volume equivalent to 4.5 embryos aliquoted into different 1.5 mL tubes before drying the metabolite extracts.

### RNA isolation

After metabolite extraction, the pellet, ceramic beads, and remaining metabolite soluble fraction (∼200 µL) of each sample was dried (∼1 h) in a Genevac EZ-2 series evaporator using low-BP lamp-off settings. Total RNA isolation was then carried out as described in detail in Pérez-Mojica *et al*., 2023^6^. In brief, 500 µL TRIzol™ (Invitrogen, USA) and 50 µL Gibco™ phosphate buffered saline pH 7.2 (Thermo Fisher Scientific, USA) were added to each sample mixing thoroughly. After 5 min incubation at room temperature, 100 µL chloroform (Sigma-Aldrich, USA) were added, samples mixed by vortex, incubated 2 min at room temperature, centrifuged for 15 minutes at 12,000 × g at 4 °C, and RNA-containing aqueous phase pipetted into a new 1.5 mL tube. RNA was precipitated by adding 250 µL ice-cold isopropanol (Sigma-Aldrich, USA) and 2 µL GlycoBlue™ (Thermo Fisher, USA), samples mixed by hand, incubated for 10 min at RT, and centrifuged for 10 minutes at 12,000 × g at 4°C. After discarding the supernatant, RNA pellets were washed with 1 ml 75% (v/v) ethanol (Pharmco, USA), air-dried, and stored at -80°C until library preparation for RNA-seq.

### Library preparation and RNA-seq

RNA-seq was carried out following our previously published step by step protocol for single-egg RNA-seq, which is based on the CEL-Seq2 method^5,6,37^. In short, RNA was resuspended in 8 µL nuclease-free water (Invitrogen, USA) and 120 nL dispensed using the I.DOT (Dispendix) into a 384-well plate holding 240 nL of primer-mix including 192 different cell barcodes with unique molecular identifiers (UMI). After first-strand (SuperScript II, Thermo Fisher Scientific) and second strand synthesis (*E. coli* DNA Pol I, Thermo Fisher Scientific) was completed, barcoded samples (n=384) were mixed into pools containing 96 samples per pool (4 libraries). cDNA was cleaned up using AMPure XP reagent (Beckman Coulter, USA) and in vitro transcription performed (MEGAscript T7 Transcription Kit, Thermo Fisher Scientific) during 16 h at 37°C. The resulting amplified RNA was fragmented using ExoSAP-IT (Thermo Fisher Scientific, USA), cleaned up using RNAClean XP (Beckman Coulter, USA), and used for first strand synthesis. Each library was diluted 1:10 (cDNA:H_2_O), an 11-cycle PCR amplification performed, and samples cleaned up using AMPure XP reagent (Beckman Coulter, USA). Custom paired-end sequencing (read 1 = 15 bp; read 2 = 250 bp) was performed using the NovaSeq 6000 instrument (Illumina) by the Genomics Core at Van Andel Institute. Sequencing depth for each individual embryo or unfertilized egg was on average 3.4 M reads, with 84% of the sequences with a Q-score ≥ 30 (FastQC version 0.11.9)^38^.

### RNA-seq data analysis, variant calling and obtaining allele specific read counts

Because the read lengths exceeded the mean fragment lengths, reads were trimmed using TrimGalore v0.6.10 with the parameters, ‘--length 15 --hardtrim5 200 --paired’. Trimmed reads were processed using a custom Snakemake workflow (https://github.com/vari-bbc/scRNAseq; commit 834077) that calls variants using a process partly adapted from GATK’s standard RNA-seq workflow (https://github.com/gatk-workflows/gatk4-rnaseq-germline-snps-indels). Reads were aligned to the BDGP6.28 (dm6) genome, from Ensembl release 100, and ERCC sequences using STARsolo from STAR v 2.7.8a^39,40^ as described step by step in Pérez-Mojica *et al*., 2023^6^. After read alignment, BAM files were split by the cell barcode tag (CB) using ‘bamtools split’ from BamTools v2.5.2^41^; the sample tag (SM) in each split BAM file was made unique by appending the CB tag via ‘samtools reheader’ from SAMtools v1.17^42^. Duplicate reads were identified based on UMIs using MarkDuplicates from PicardTools v3.0.0 (http://broadinstitute.github.io/picard/) with the parameters, ‘--BARCODE_TAG “UB” --VALIDATION_STRINGENCY SILENT’. For non allele-specific analysis, samples with a total raw read count <150,000 or transcripts with <5 read counts on <5 samples were filtered out from the analysis. Read count normalization, unsupervised sample clustering, transcriptome entropy calculi, generation of a lineage tree, identification of unfertilized eggs and pseudo-temporal ordering of samples was carried out using R package RaceID v0.3.0^11^ as previously described^5,6^. For allele-specific analysis, variants were called using the joint genotyping workflow in GATK v4.4.0.0^43^ as suggested in Brouard et al., 2019^44^. Reads with Ns in the CIGAR string were split using ‘SplitNCigarReads’. Base qualities were recalibrated using ‘BaseRecalibrator’ and ‘ApplyBQSR’ using the DGRP2 variants^7^ (dm6 liftover downloaded from https://www.hgsc.bcm.edu/arthropods/drosophila-genetic-reference-panel on Mar 30, 2021) for the set of known variants. ‘HaplotypeCaller’ was run with the parameters, ‘-ERC GVCF -dont-use-soft-clipped-bases --standard-min-confidence-threshold-for-calling 20’, on each embryo (cell barcode), for each chromosome separately to reduce runtime. Joint genotyping was conducted on each chromosome separately, by running ‘GenomicsDBImport’ then ‘GenotypeGVCFs’. Per-chromosome VCF files were sorted using ‘SortVcf’ then merged into a single VCF using ‘MergeVcfs’. Variants were filtered by running ‘VariantFiltration’ with the parameters, ‘--window 35 --cluster 3 --filter-name "FS" --filter "FS > 30.0" --filter-name "QD" --filter "QD < 2.0" --genotype-filter-name "GQ" --genotype-filter-expression "GQ < 15.0" --genotype-filter-name "DP" --genotype-filter-expression "DP < 10.0"’, followed by ‘SelectVariants’ with the parameters, ‘--exclude-filtered --set-filtered-gt-to-nocall’. Genic variants were annotated using ‘eff’ from SNPEff v5.1^45^, with the parameters, ‘-no-downstream -no-intergenic -no-upstream’, and the pre-configured ‘BDGP6.32.105’ database. A table containing variant coordinates, variant information and allelic depth was extracted from the VCF file using ‘VariantsToTable’ from GATK v4.4.0.0 using the parameters, ‘-F CHROM -F POS -F REF -F ALT -F TYPE -F ANN -GF AD’.

Downstream analyses were conducted in R v4.2.1 and the script can be found at https://github.com/vari-bbc/scRNAseq/blob/main/scripts/ASE_example.Rmd. The allelic depth table was imported as a GenomicRanges object (v1.50.2)^46^. Only biallelic SNPs annotated to a single gene on standard chromosomes or “mitochondrion_genome” were retained. A SummarizedExperiment object was created to store the variant coordinates, the sample metadata, and the REF and ALT allelic depths as separate assays. Variants were further filtered to retain only those with at least 5 samples with > 5 read counts for each of REF and ALT (not necessarily the same samples); for mitochondrial variants, if either REF or ALT allelic depths passed the above filter, the variant was retained. The DGRP2 VCF file was imported and processed using the VariantAnnotation v1.44.1 package^47^, retaining only polymorphic variants between line_737 and line_352. DGRP2 variant information was merged with the SummarizedExperiment based on matching chromosomal coordinates, and reference and alternate alleles. Where there was a matching entry, parental origins for alleles were determined based on the parental genotypes identified in DGRP2 and used to derive maternal and paternal counts, which were stored as additional assays in the SummarizedExperiment object. Variants annotated to the same gene were ranked in descending order based on the number of samples expressing both alleles (> 0 counts), and the total counts (reference or alternate) across samples. Diagnostic plots of the maternal and paternal counts were created by coercing the SummarizedExperiment object into a SingleCellExperiment object (v1.20.1)^48^ and utilizing functions from scater v1.26.1^49^ on only the top-ranked variant of each gene. We use FlyBase (release FB2024_01) to find information on gene function and gene expression.

### Weighted gene correlation network analysis (WGCNA)

To reduce the complexity of the transcriptomic data in both allele-specific and non-allele specific datasets, we employed weighted gene coexpression network analysis via the R v4.3.0 (https://cran.r-project.org/) package *WGCNA*^15^ following a workflow similar to https://pklab.med.harvard.edu/scw2014/WGCNA.html. Before the WGCNA was run, any genes with at least 2 unique paternally deposited reads in any unfertilized embryo were removed from the paternally deposited data. Then the unfertilized eggs were removed from all datasets and pseudo-times were refactored to maintain consecutive integer pseudo-times. Finally, to prevent the allele-specific data from being too sparse, which can impact WGCNA fits, only genes with at least 2 unique reads in 2 or more embryos were kept. Because we are investigating development patterns, the direction of expression (upregulated or downregulated at a given pseudo-time) mattered. Therefore, all WGCNA networks were created using a signed-hybrid correlation. The soft power threshold for each network was chosen based on a combination of R^2^ exceeding 0.8 and having mean connectivity in the transition from high to low (visually this is a soft-power in the ‘elbow’ of the mean connectivity plot). For non allele-specific data we chose a soft-power of 6, for paternally deposited transcripts 12, and maternally deposited 7. The minimum module size for non allele-specific data was set to 20 to allow for any developmental patterns summarized by a small subset of genes, and for the allele-specific data the minimum module size was reduced to 10 due to these datasets having far fewer total genes. All modules that were highly correlated (R > 0.95) were merged into one. Pearson correlations were then used to determine which metabolites are associated with specific module eigengenes. To account for multiple testing, p-values were adjusted using Benjamini-Hochberg to control the false discovery rate at 5%. T A correlation network built on all pairwise module:metabolite correlations was created using the R package i*graph*^50^ and then exported to Cytoscape (https://cytoscape.org/) via *Rcy3*^51^ for additional visualization. Code for this workflow is available on Github (https://github.com/vari-bbc/Drosophilia_Single_Embryo_Workflow.

### Activation clusters

To estimate the exact transcriptional onset for each maternally and paternally deposited gene, generalized additive models (GAM) were fit individually to the RUVg normalized allele-specific expression data via the R v4.3.0 (https://cran.r-project.org/) package *mgcv*^52^. GAMs were fit using the restricted maximum likelihood method and shrunken cubic splines as the smoothing term. A given gene’s activation time was set at the first pseudo-time where the lower bound of the 95% confidence interval exceeded 1 unique read. The activation times for each gene were then passed to the function *select* within the R package *mixR*^53^ to determine if the joint distribution of activation times had multiple modes. All four possible distribution families were run (normal, log-normal, gamma, and Weibull) with modes ranging from 1 to 6 and both equal and unequal variances were examined. The mixture fit with the lowest Bayesian information criterion was considered the best model and was then used to group each gene into activation clusters. For the paternally deposited data, the best fit was a log-normal mixture, with unequal variances, and 4 modes. The function *mixfit* from *mixR* was used to fit the mixture model and visualize the results. Code for this analysis is available on Github in the WGCNA workflow (https://github.com/vari-bbc/Drosophilia_Single_Embryo_Workflow).

### Metabolite identification

In preparation for metabolite identification, samples were resuspended in 30 µL water (90 µL for QCs) (Optima™ LC/MS Grade Fisher Chemical™), pulse vortexed, sonicated for 5 min at 40 kHz in a Bransonic® bath sonicator (Branson Ultrasonics, USA), centrifuged for 3 minutes at 17,000 × g at 4°C, and transferred to 0.3 mL clear glass vials with fused-in inserts (Supelco, Millipore Sigma, USA). Samples were first run using an ion-paired method, completely dried in a Genevac EZ-2 series evaporator (ATS Life Sciences Scientific Products) using low-BP lamp-off settings, and stored at -80°C until later performing a non-ion-paired method. Both methods were based on ultra-high performance liquid chromatography coupled to a 6470 triple quadrupole mass spectrometer (Agilent Technologies, USA). For the ion-paired we used column ZORBAX RRHD Extend-C18, 80Å, 2.1 x 150 mm, 1.8 µm, 1200 bar pressure limit (Cat # 759700-902 from Agilent), solvent A (H2O with 10mM Tributylamine 15mM Acetic Acid, and 0.1% Medronic Acid), and solvent C (Methanol with 10mM Tributylamine 15mM Acetic Acid, and 0.1% Medronic Acid). For the non-ion-paired we used column CORTECS T3 Column, 120Å, 1.6 μm, 2.1 mm X 150 mm, 1/pk (Cat # 186008500 from Waters), solvent A (H2O with 0.1% formic acid) and solvent B (90% Acetonitrile with 0.1% formic acid). We also use 99% Acetonitrile to recondition the column between runs. Randomization was performed ensuring that a similar number of embryos and unfertilized eggs of all developmental time windows were allocated to a specific run batch. In each run, the instrument performance on a solvent blank, a metabolite standard mix and two QCs injections was evaluated before starting to inject the same QC every 8-11 samples (a new QC aliquot per batch). After metabolite determination on individual samples was performed, remaining metabolite extracts (stored at -80°C) were resuspended, mixed according to their pseudo-time (10 embryos per pool), dried, prepared for metabolite identification as described above, and re-run using the same ion-paired and non-ion-paired methods. Since analysis of metabolites in male and female embryos did not reveal any significant sex-specific differences we disregarded sex for sample pooling. The LC and MS parameters for both methods and transitions for our screening panel of 155 metabolites can be found in **Supplementary Methods1**.

### Metabolite data processing and analysis

Synchronized peak integration was performed on all individual embryos, unfertilized eggs, QCs, and blanks from all batches using Skyline v23.1.0.268^54^. Metabolite peak areas (raw data) were corrected for instrument performance (within-batch correction) and batch differences (between-batch correction) using QC-based random forest signal correction, QCspan=0.75, and coCV=20 (statTarget v1.28.0 R package)^55^. This QC-correction was only applied to metabolites that showed a decrease in variance upon correction, otherwise raw values were scaled making the average QC signal for a specific metabolite the same in all batches (between-batch correction). Outlier detection was performed based on 99.7% prediction intervals for General Additive Models (GAM) on the embryo metabolite values aligned according to the RNA-seq pseudo-time trajectory (n=245). To facilitate analysis, metabolite data was divided by the average value of each metabolite in embryos (mean normalization), making normalized data to average 1 for each metabolite. A similar data processing was carried out for pooled samples except for outlier identification using GAM.

### Sex-specific analysis of gene expression and metabolite abundance

RNA-seq normalized read counts of each transcript or normalized abundance of each metabolite were compared between male and female embryos using splineTimeR v1.26.0^56^. Each embryo with known sex was considered as a replicate in every RNA-seq cluster from RaceID analysis (time points). To mitigate potential bias from the start and end points of the pseudo-time on the analysis, potentially driven by development rather than sex, we generated additional clusters at the beginning and end to bias the first and last time point towards the null. For the first cluster we used 10 embryos prior sex identification and assigned them as male or female. For the last cluster, we duplicated the last 10 embryos in the pseudo-time and assigned them as male or female. Transcripts with <3 reads in <10 samples were excluded from analysis. Significance for both transcripts and metabolites was established at a Benjamini-Hochberg adjusted p-value (padj) <0.05 using three degrees of freedom.

### *Rnrs* and *dnk* RNAi knockdown

To induce the RNAi- knock-down (KD), females carrying a Gal4-driver (BDSC_7063) were crossed with males carrying a transgene encoding the shRNA (BDSC_44022 for RnrS, BDSC_65886 for dnk). The resulting offspring (F1), which are producing oocytes that are KD for the target gene, were then crossed with each other, and the pupae of the second generation (F2) were counted for 5 independent crosses. This approach allows KD in late-stage ovaries without interfering with meiosis, or fertilization as previously described^33^. For control crosses, we mated females carrying the Gal4-driver with males carrying an empty attP40 landing site (VDRC_60100).

### DATA ACCESS

● Raw sequencing files and processed data files from this study have been deposited at NCBI Gene Expression Omnibus (GEO) under accession number GSE263568 and will be released after publication in a peer reviewed journal (temporal token for reviewers was provided to the editor).
● Normalized metabolite data of all unfertilized eggs and embryos included in pseudo-time analysis can be found Table S5. Scaled metabolite data for embryos according to developmental trajectory (pseudo-time) can be found in Table S6.
● All code used is available on our Github repository at https://github.com/LempradlLab/Drosophila_embryo_metabolism/.
● Any additional information required to reanalyze the data reported in this paper is available from the lead contact upon request.

## Supporting information

all_supplemental_files

## ACKNOWLEDGEMENTS

We thank the Van Andel Institute Genomics Core (RRID:SCR_022913), specially Marc Wegener and Marie Adams, for their assistance with RNA-seq. We thank the Van Andel Institute Bioinformatics and Biostatistics Core (RRID:SCR_024762) for supporting the analysis. We thank the Van Andel Institute Mass Spectrometry Core (RRID:SCR_024903) for their assistance in metabolomic analysis. Figure 1A was created using <u>BioRender.com</u>. Stocks obtained from the Bloomington Drosophila Stock Center (NIH P40OD018537) were used in this study. This research was funded by the Van Andel Institute.

## AUTHOR CONTRIBUTIONS

J.E.P-M., R.D.S., and A.L. designed and directed the study. J.E.P-M., C.N.I., and J.R. performed experiments. J.E.P-M., Z.B.M., K.H.L. and A.L. analyzed the data. K.H.L. created workflows for non-allele-specific and allele-specific gene expression analysis. Z.B.M. created workflows for WGCNA and activation clusters. J.E.P-M., and A.L. wrote the original draft. R.D.S., and Z.B.M. reviewed and edited the manuscript.

---

**Figure S1.**
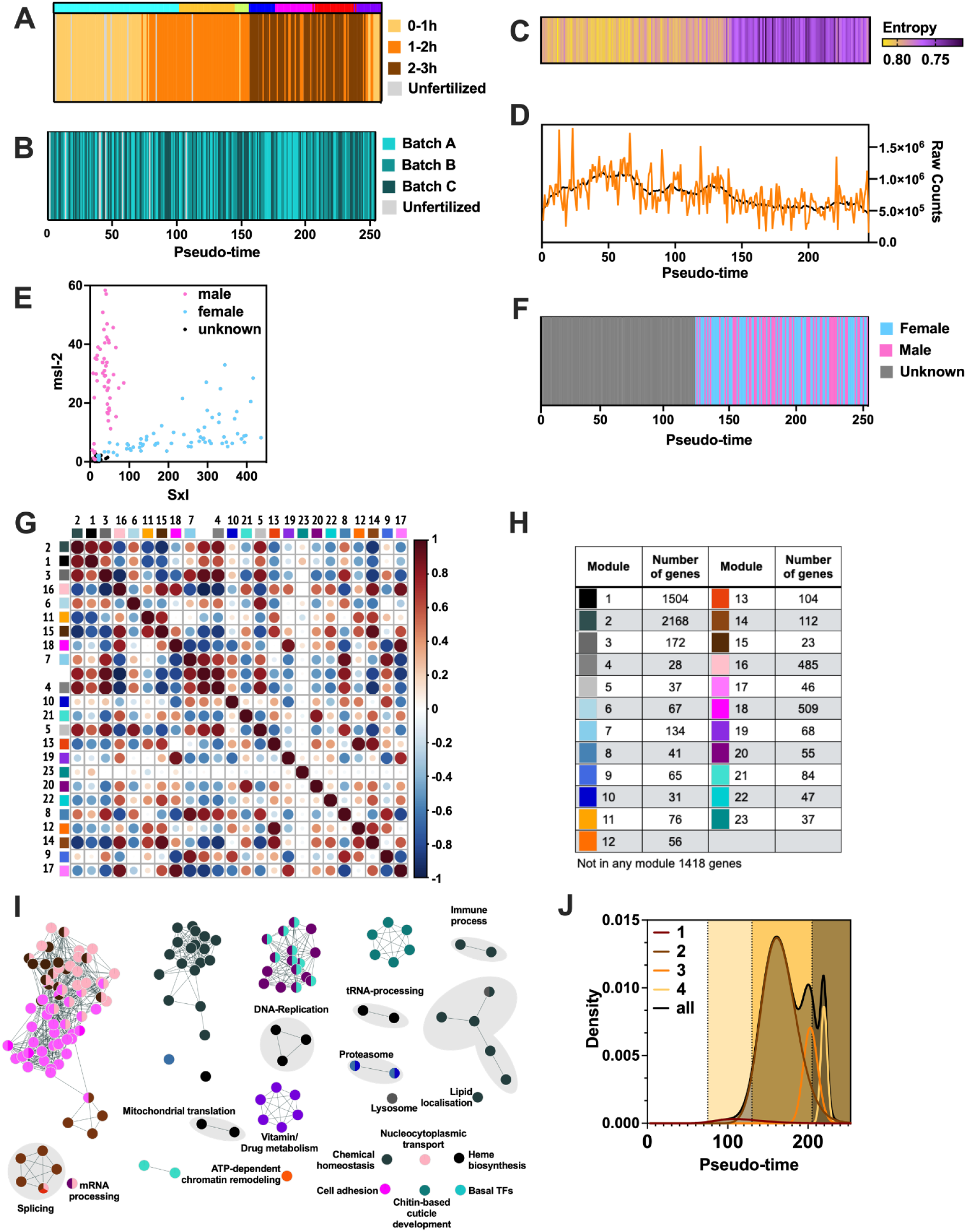
Bioinformatic analyses using the early embryo transcriptome. Comparison of the pseudo-time order indicating **(A)** the actual collection time intervals for embryos and unfertilized eggs, colors in the top bar indicate clusters from Figure 1B; **(B)** the mating cage; (**C**) the transcriptome entropy (**D**) the total number RNA-seq raw read counts per embryo. **(E)** XY Plot of Sex lethal (Sxl) expression on the x-axis and male-specific lethal 2 (msl-2) expression on the y-axis of all samples included in the pseudo-time. **(F)** Embryos in pseudo-time order indicating the sex for each sample. **(G)** Correlation heatmap of the identified WGCNA modules. **(H)** Table showing color code and total number of genes in each WGCNA module. **(I)** Network map of overrepresented pathways in modules, pathways are linked according to gene-set overlap. Overrepresentation analysis using g:profiler (significance threshold g:SCS=0.1) was used as input into Cytoscape for network visualization. **(J)** Distribution of zygotic mRNAs in activation time windows indicated by numbers.

**Figure S2.**
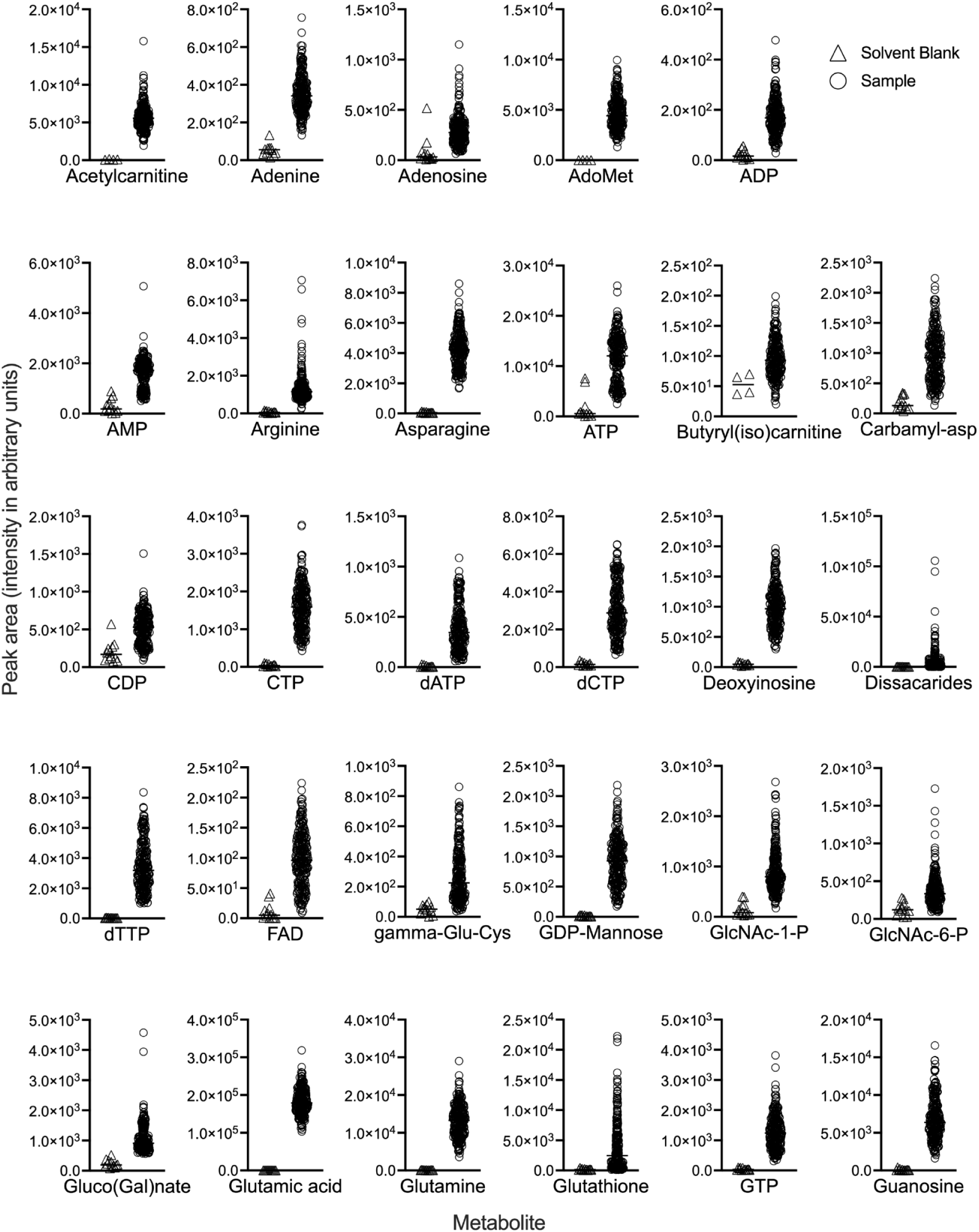
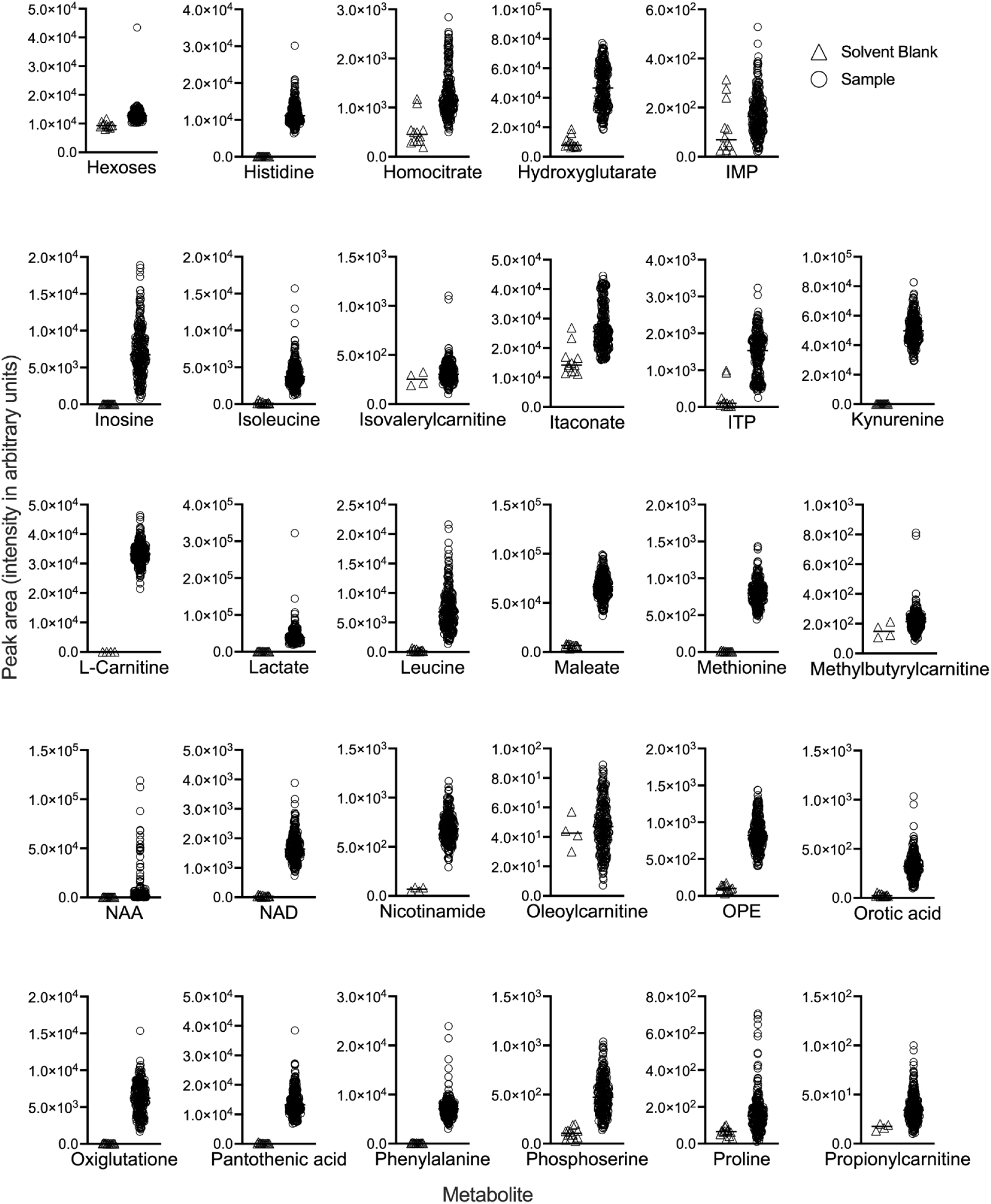
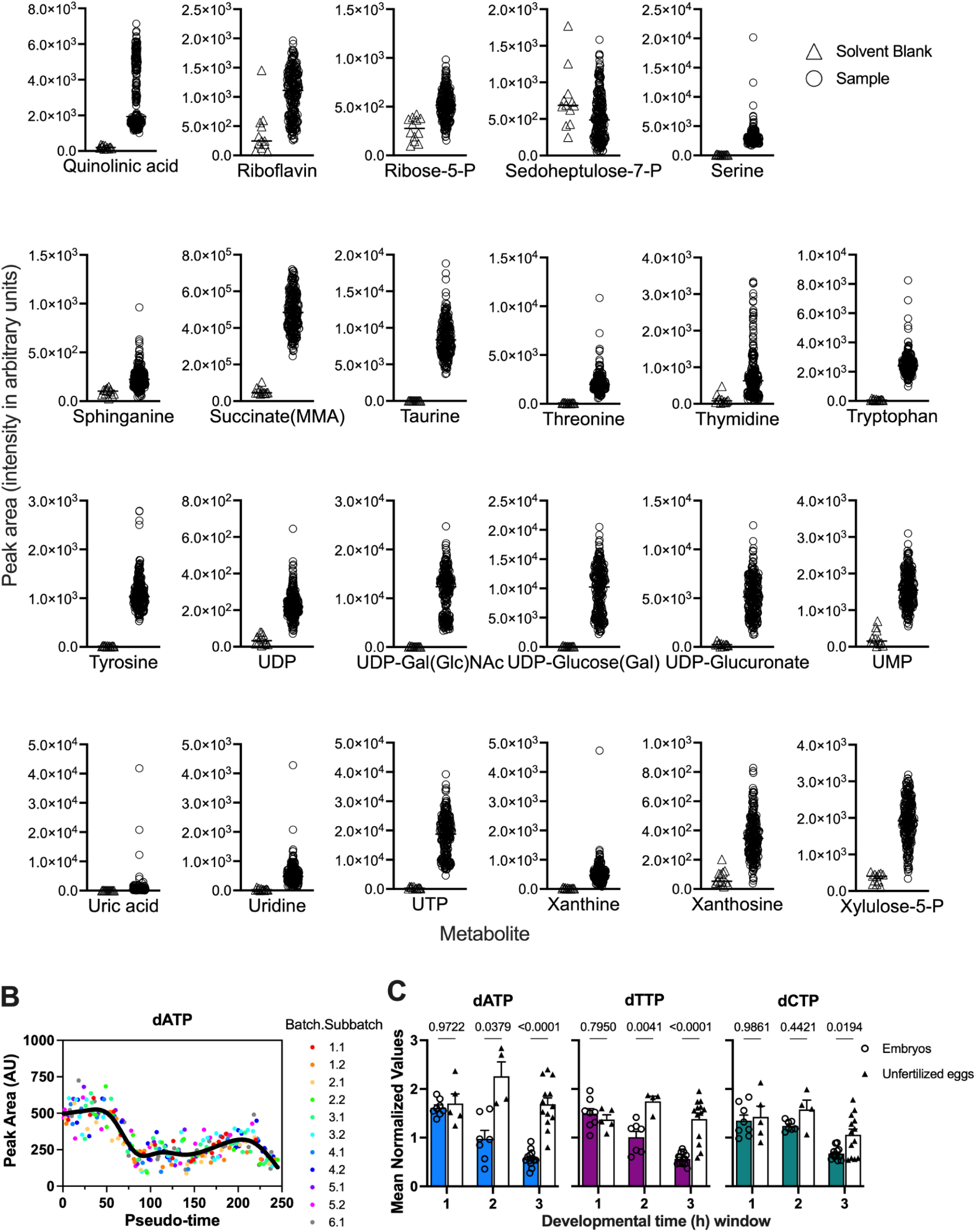
Single-embryo detection of polar metabolites. **(A)** Metabolite abundance in arbitrary units (AU) for all 81 metabolites identified. **(B)** dATP abundance plotted along the RNA-seq pseudo-time order. Each sample is colored according to the LC-MS run batch. **(C)** Comparison of dATP, dTTP, and dCTP relative abundance between embryos (colored bars) and unfertilized eggs (open bars) during the first 3 hours of development.

**Figure S3.**
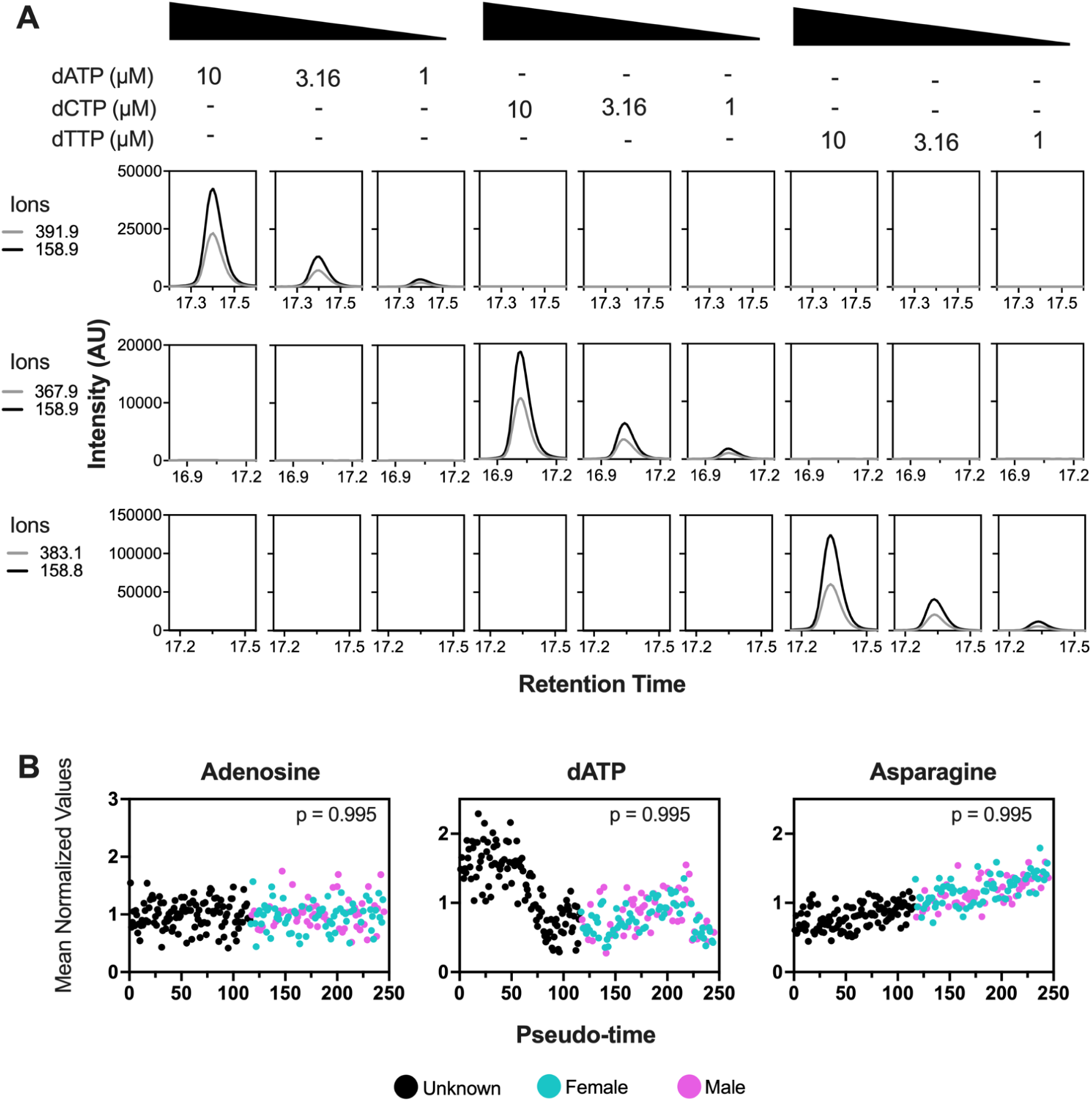
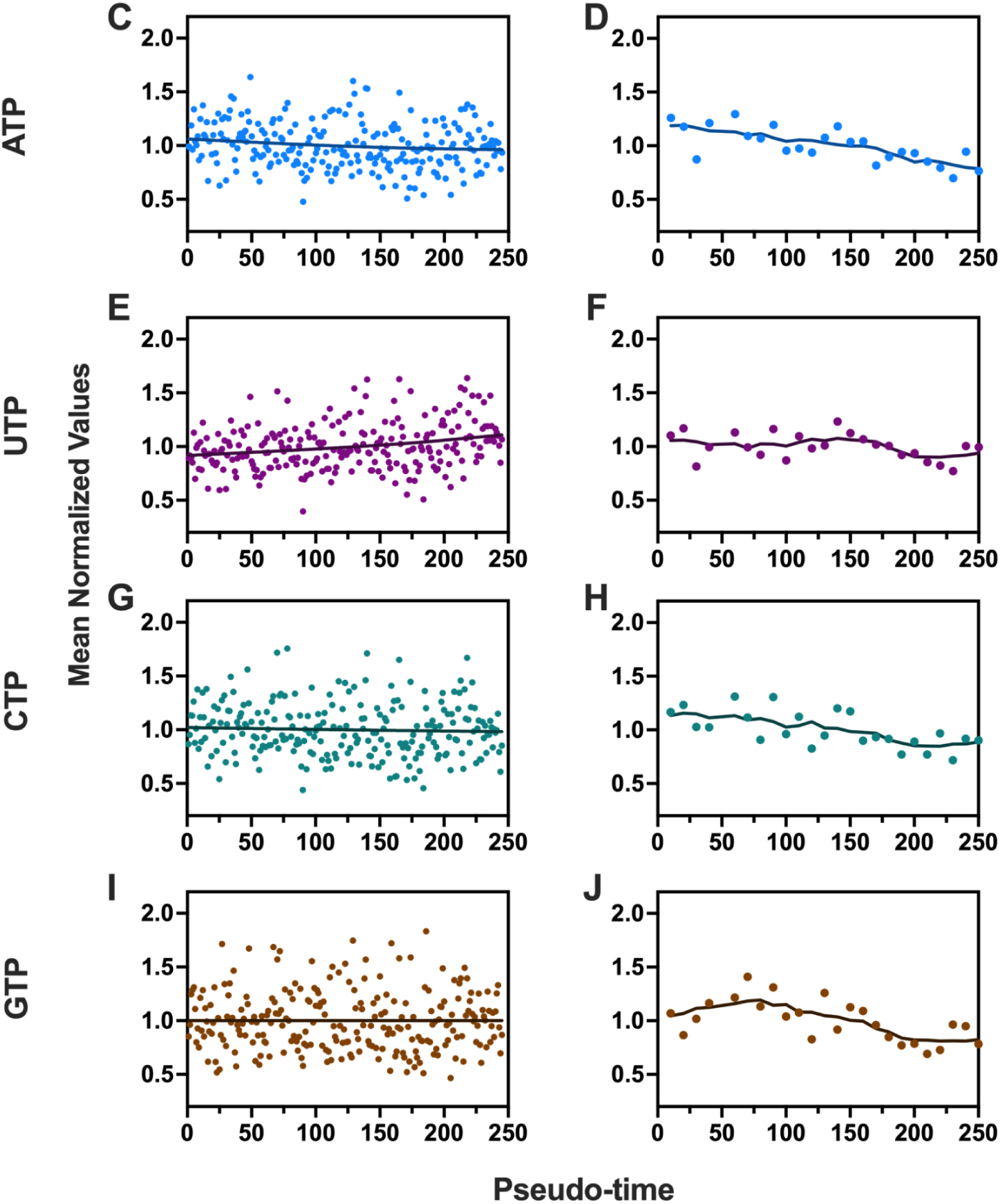
Validation of dNTPs transitions, and sex and nucleotides metabolite patterns. **(A)** Ion chromatograms for transitions of dATP, dTTP and dCTP in commercial standards (Thermo Fisher, USA); two transitions of dATP (490➝391.9, 490➝158.9), two transitions of dTTP (481➝383.1, 481➝158.8), and two transitions of dCTP (466➝367.9, 490➝158.9) at different concentrations. **(B)** Examples for relative metabolite abundance in male and female embryos. Embryos in pseudo-time order colored according to the sex for each sample and P-value from splineTime R analysis. **(C-J)** Nucleotide relative abundance in embryos according to pseudo-time order (C, E, G, I) by single-embryo LC-MS or (D, F, H, J) in pooled samples (n=10 embryos/each).

**Figure S4.**
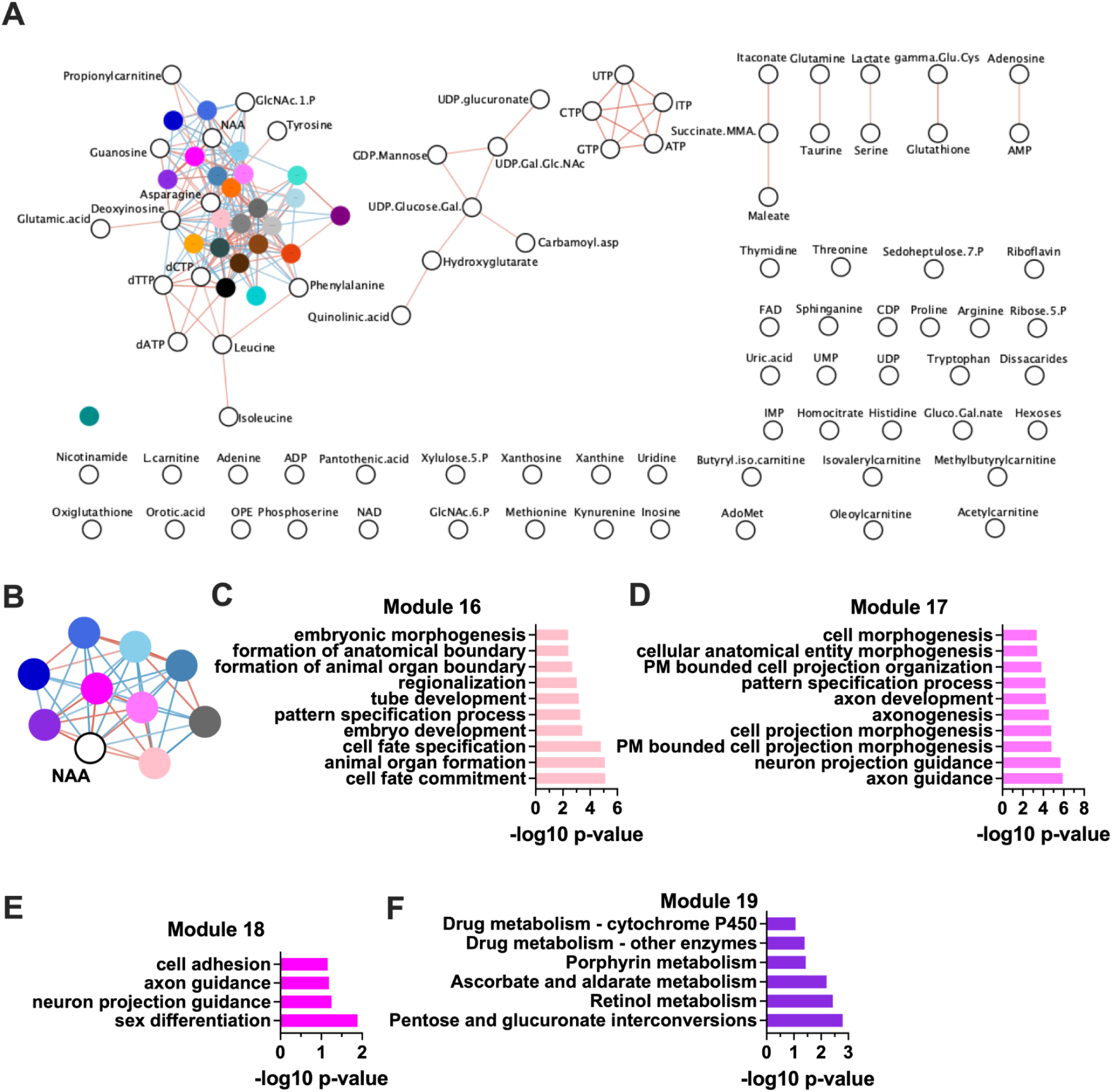
Relationship between metabolic and transcriptional patterns in the early Drosophila embryo. **(A)** Network map of correlations between WCGNA transcript modules (colored circles) and metabolites (white circles) and **(B)** Network map of N-Acetyl Aspartate (NAA) and its neighbors using network visualization with Cytoscape. Red or blue connecting lines indicate positive or negative correlations, respectively. **(C-F)** Overrepresentation analysis for transcript in modules **C)** 16, **D)** 17, **E**)18, and **F)** 19 using g:profiler (significance threshold g:SCS=0.1).

